# Defective Integrator activity shapes the transcriptome of patients with multiple sclerosis

**DOI:** 10.1101/2023.11.24.568591

**Authors:** Yevhenia Porozhan, Mikkel Carstensen, Sandrine Thouroude, Mickael Costallat, Christophe Rachez, Eric Batsché, Thor Petersen, Tove Christensen, Christian Muchardt

**Author notes:** Correspondance : Christian Muchardt, ORCID : https://orcid.org/0000-0003-0145-4023, Street address : CNRS – UMR8256 – Biological Adaptation and Ageing Institut de Biologie Paris-Seine - FR3631, Sciences Sorbonne Université Campus Pierre et Marie Curie, 7-9, Quai Saint Bernard - Bât A, 2ème étage 75252 Paris Cedex 05, France. Participated equally to this work (Yevhenia Porozhan, Mikkel Carstensen). Participated equally to this work (Thor Petersen, Tove Christensen).

## Abstract

HP1α/CBX5 is an epigenetic regulator with a suspected role in multiple sclerosis (MS). Here, using high-depth RNA sequencing on monocytes, we identified a subset of MS patients with reduced CBX5 expression, correlating with progressive stages of the disease and extensive transcriptomic alterations. Examination of rare non-coding RNA species in these patients revealed impaired maturation/degradation of U snRNAs and enhancer-RNAs, indicative of a reduced activity of the Integrator, a complex with suspected links to increased MS risk. At protein-coding genes, compromised Integrator activity manifested in reduced pre-mRNA splicing efficiency and altered expression of genes regulated by RNA polymerase II pause-release. Inactivation of Cbx5 in the mouse mirrored most of these transcriptional defects and resulted in hypersensitivity to Experimental Autoimmune Encephalomyelitis (EAE). Collectively, our observations suggested a major contribution of the Integrator complex in safeguarding against transcriptional anomalies characteristic of MS, with HP1α/CBX5 emerging as an unexpected regulator of this complex’s activity. These findings bring novel insights into the transcriptional aspects of MS and provide potential new criteria in patient stratification.

---

Multiple sclerosis (MS) is an acquired demyelinating disorder of the central nervous system (CNS) characterized by chronic inflammation and the formation of scar tissue (sclerosis) in various areas of the CNS. It is the most common disabling neurological disease of young adults, and currently available treatments primarily focus on managing the symptoms.

During the course of the disease, immune cells breach the blood-brain barrier (BBB) and attack the myelin sheath that insulates neurons. This leads to symptoms like numbness in various body parts, paresis, coordination and balance difficulties, blurred vision, slurred speech, and cognitive changes. MS can manifest in several distinct forms. Clinically Isolated Syndrome (CIS) represents the initial clinical manifestation of a condition with features of inflammatory demyelination, which may suggest the possibility of MS, but does not yet fulfill the criteria for dissemination in time and space required for a definitive diagnosis. Once criteria for dissemination in time are met, the most common form of MS is Relapsing-Remitting MS (RRMS), characterized by clearly defined relapses of symptoms followed by periods of partial or complete recovery. RRMS may evolve into Secondary Progressive MS (SPMS), where the disease worsens gradually without distinct relapses or remissions. Another form, Primary Progressive MS (PPMS), is characterized by a gradual worsening of symptoms from the onset, without distinct relapses or remissions (1).

The etiology of the disease remains unclear and is a subject of ongoing debate. Generally, it is believed that CNS-directed autoimmunity in MS results from a complex interplay of genetic susceptibility, hormonal factors, and environmental cues. These environmental cues, which encompass lifestyle aspects, include Epstein-Barr virus (EBV) infection, exposure to tobacco smoke and organic solvents, obesity during adolescence, and limited sun exposure coupled with low vitamin D levels. Some of these factors, particularly EBV serology, obesity, and possibly vitamin D deficiency, seem to be particularly relevant when they occur during adolescence (2). This early exposure contrasts with the typical onset of multiple sclerosis, which often does not manifest until the second or third decade of life, hinting at the existence of a long-term memory mechanism within the immune system. While the modulation of the peripheral adaptive immune response has been extensively studied in this regard (3), the role of epigenetic alterations in creating a lasting imprint on the immune cells’ transcriptional activity represents an underexplored but potentially significant memory mechanism, bridging the gap between early-life environmental exposures and later disease onset.

Previously, we have explored this possibility by examining the activity of HP1α/CBX5, a transcriptional repressor with affinity for both chromatin and RNA (4–6), and encoded by a gene sharing one of its promoters with hnRNPA1, a gene harboring SNPs previously associated with elevated risk of MS (7). While CBX5 is best known for its affinity for the heterochromatin-enriched histone H3 lysine 9 tri-methylation (H3K9me3) mark, it is also active in euchromatin (8,9). Accordingly, we showed that CBX5 was involved in repressing a set of cytokine genes and that in patients with MS, its recruitment to the promoters of these genes was reduced. This reduced recruitment could at least in part be correlate with increased citrullination of histone H3 at arginine 8 (H3Cit8), destroying the CBX5 binding site on the histone tail (10). Thus, reduced CBX5 activity in patients with MS offers a potential rationale for the concurrent elevation in the activity of proinflammatory cytokine genes and normally-heterochromatinized DNA repeats, such as endogenous retroviruses (HERVs).

In a more recent study, we have further shown that at least a fraction of the HERV transcripts produced in MS patients are enhancer RNAs (eRNAs) originating from viral sequences coopted as immune-gene enhancers, and reactivated in the patients (11). Thus, CBX5 may also function as a regulator of eRNA production. Aligning with such a function, other HP1 family proteins are known to participate in RNA metabolism; notably, SWI6 is responsible for the degradation of heterochromatic transcripts in yeast (12), and HP1γ (CBX3) influences the regulation of alternative splicing in mammals (13–17). We therefore wished to re-explore the role of CBX5 in MS in the light of high-depth genome-wide RNA-seq data allowing for detection of rare RNA species such as eRNAs and alternative splice junctions. This analysis has revealed that reduced levels of HP1α/CBX5 in monocytes from MS patients correlate with a range of transcriptional abnormalities all indicative of diminished Integrator complex (INTcom) activity. Thereby, the INTcom, crucial for non-coding RNA (ncRNA) maturation and RNA Polymerase II (RNAPII) elongation, emerges as a novel player in MS pathology, offering mechanistic insights into numerous gene deregulation events associated with the disease.

## RESULTS

### A subset of MS patients displays extensive transcriptional deregulation in monocytes

With the objective of obtaining high quality RNA-seq allowing detection of rare RNA species, we collected monocytes from a cohort of 18 MS patients and 7 control patients (Figure 1A). To explore eventual changes in the RNA populations induced by the disease progression, we selected a very heterogeneous set including patients with familial MS (#68, #125, and #130 – one family, 3 generations), under immune modulating treatment (#125, #127, #135, #136, and #139), or with comorbidities (psoriasis, IDDM, Graves, Crohn and ulcerative colitis). The control group, designated as symptomatic controls (SCs), comprised individuals who had sought medical assistance for symptoms suggestive of MS, but who, upon examination, exhibited no objective clinical or paraclinical findings conclusive for a specific neurological disease at the time of sample collection (Figure 1A). The RNA-seq analysis was conducted using a stranded paired-end protocol, yielding an output of approximately 110 million reads per sample. Principal Component Analysis (PCA) on the RNA-seq data, profiling the 500 most variable genes, identified a substantial proportion of variance in gene expression across the samples. It further revealed the existence of two distinct biological subgroups, with PC1 scores that were either lower or significantly higher than those of the SCs (Figure 1B and Sup. Tables 1A, 1B, and 1C).

**Figure 1:**
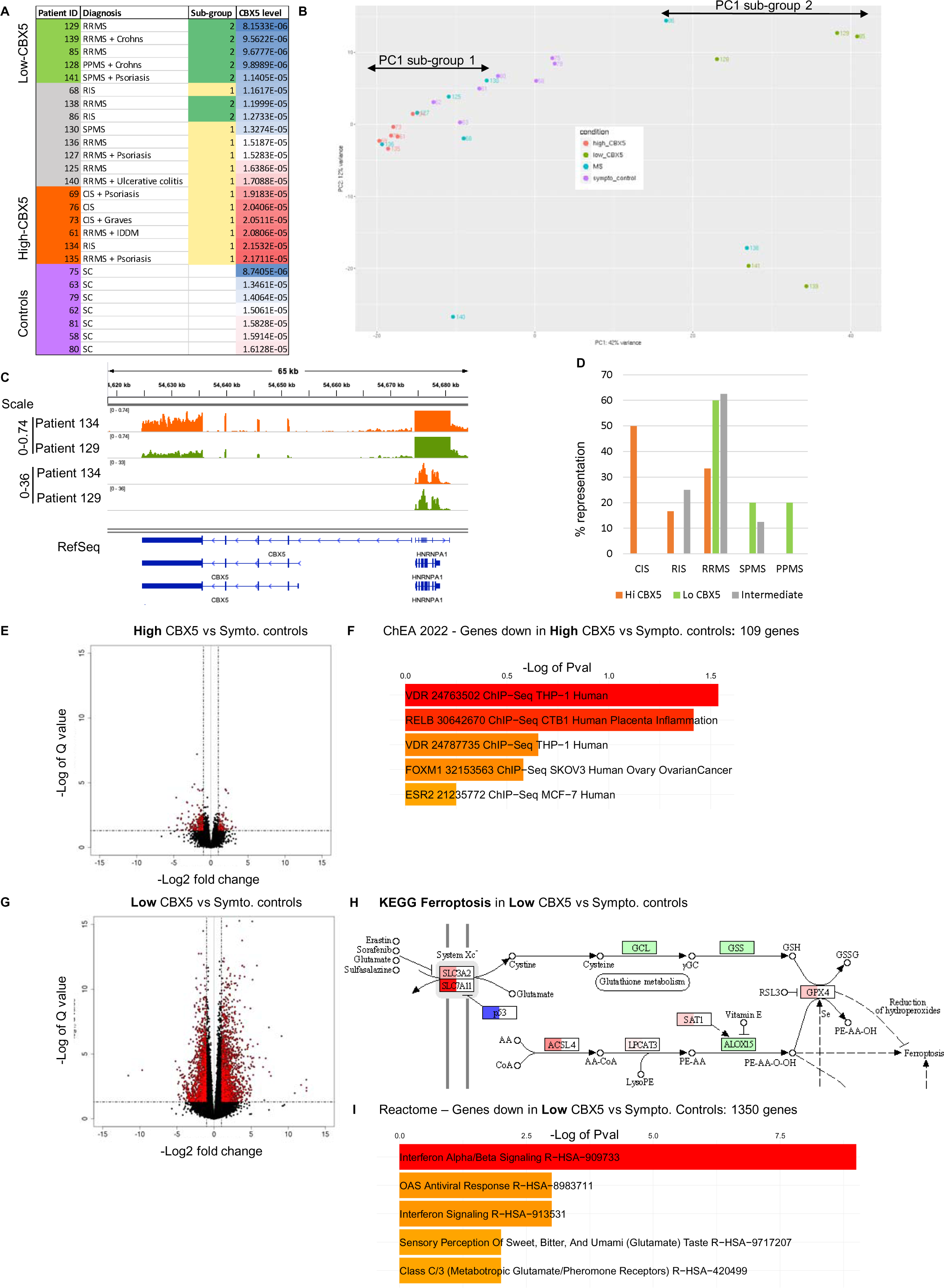
Extensive transcriptional deregulation in monocytes from patients with reduced expression of CBX5. (A) List of patients with MS with indications on the stage of the disease and eventual comorbidities. Symptomatic controls (SC) are also indicated. Donors are ranked according to levels of CBX5 expression after normalization by DESeq2. Patients highlighted in red express CBX5 at high levels (upper quartile), while patients indicated in green express this gene at low levels (lower quartile). (B) PCA analysis using the 500 most variable genes. (C) Graphic representation of the CBX5-HNRNPA1 locus using the IGV genome browser. Bar charts represent the normalized expression levels of the two genes in the indicated patients. The 2 top lanes use a scale suited for the visualization of CBX5, while the 2 following lanes are at a scale fitted for HNRNPA1. RefSeq allows visualization of splice variants of the two genes annotated in Hg19 version of the human genome. (D) Within each group of patients defined by either high, low, or intermediate CBX5 levels, the percentage representation of each stage of the MS disease is indicated. (E) Vulcano plot illustrating differential gene expression between Hi-CBX5 patients and symptomatic controls. (F) Analysis of genes down-regulated between Hi-CBX5 patients as compared symptomatic controls to using the ChEA (ChIP-Seq Enrichment Analysis) database providing information on transcription factors likely to regulate these genes. (G) Vulcano plot illustrating differential gene expression between Lo-CBX5 patients and symptomatic controls. (H) Schematic representation of the KEGG ferroptosis pathway. Genes up- or down-regulated between symptomatic controls and Lo-CBX5 are indicated in red or blue respectively. (I) Analysis of genes down-regulated in Lo-BX5 compared to symptomatic controls using the Reactome pathway databank revealing functional enrichment.

We further noted that expressions levels of CBX5 varied among patients and that ranking the patients according to their respective CBX5 expression level essentially recapitulated the PCA. Indeed, patients with CBX5 expression levels in the lower quartile (Lo-CBX5) largely coincided with the patients strongly affected in their transcriptome, while patients with CBX5 expression levels in the upper quartile (Hi-CBX5) remained transcriptionally close to the SCs (Figure 1A, 1B, and 1C). The highly transcriptionally divergent/Lo-CBX5 patients were all either RRMS, SPMS, or PPMS, while the Hi-CBX5 group was enriched in early-stage patients (Figure 1D). None of the other criteria proved relevant (patient age, gender, or comorbidities).

Pathway analysis on the genes differentially expressed between Hi-CBX5 patients and SCs (38 upregulated genes and 109 down-regulated genes, 2-fold or more, adj. pVal<0.05, baseMean>10 – Figure 1E) revealed that down-regulated genes were enriched in Vitamin D regulated genes (analysis of the ChEA database, Figure 1F). This observation may reflect the importance of Vitamin D in early phases of the disease.

In contrast to the Hi-CBX5 patients, Lo-CBX5 patients displayed a dramatically high number of genes with modified expression when compared to the SCs, with a total of 1395 upregulated and 1350 down-regulated genes (2-fold or more, adj. pVal<0.05, baseMean>10 – Figure 1G). As anticipated from earlier studies, pathway analysis using the KEGG database on upregulated genes identified activation of the IL17-, TGF-beta-, NFkappaB-, and MAPK-pathways as well as genes associated with hypoxia (18) (Sup. Figure 1A). The pathway analysis also detected a significant enrichment in genes associated with ferroptosis. This form of regulated cell death characterized by the iron-dependent accumulation of lipid peroxides, has previously been linked to neurodegeneration in MS (19). It is induced either through inactivation of the glutathione peroxidase GPX4, the enzyme responsible for detoxifying lipid hydroperoxides, or by functional inhibition of the cystine-glutamate antiporter, a two-subunit complex composed of SLC7A11 and SLC3A2. Yet, in the monocytes of the Lo-CBX5 patients, while p53 (TP53), which exerts a protective role against ferroptosis was strongly down-regulated (2-fold, Adj. pVal<10^-4^), we observed only a moderate upregulation of enzymes promoting lipid peroxidation (ACSL4 and LPCAT3). In addition, the detoxifying pathways (SLC7A11, SLC3A2, and GPX4) were markedly upregulated, and markers of ferroptosis were either down-regulated PTGS2 or unchanged (CHAC1) (Figure 1H and Sup. Figure 1B). Although transcriptional activity is only indicative on enzymatic activities, this suggests that pro-ferroptotic signals are activated in the Lo-CBX5 monocytes, but are offset by compensatory mechanisms inhibiting cell death.

Analysis of the down-regulated genes with either Reactome, BioPlanet, or WikiPathway databases revealed a significant enrichment of genes associated with Type I interferon signaling (Figure 1I and Sup. Figure 1C). This is consistent with the observed benefits of administering interferon alpha/beta to patients with MS (20).

In conclusion, most of the traits commonly linked with individuals with MS, such as heightened activation of stress pathways and diminished activity of type I Interferon, are predominantly observed in patients who exhibit lower levels of CBX5 expression, in association with a significantly disrupted transcriptome. In contrast, patients expressing high levels of CBX5, enriched in patients in early phases of the disease, display only a minor deregulation of their transcriptome, exhibiting reduced activity of vitamin D reactive genes as their main characteristic.

### Evidence for reduced Integrator activity in Lo-CBX5 patients

To investigate the regulatory mechanism underlying the extensive transcriptome reprogramming observed in Lo-CBX5 patients, we confronted the list of genes upregulated in Lo-CBX5 patients (relative to the SCs) with the ENCODE Transcription Factor Data Bank, which serves as a comprehensive repository of information about transcription factors and their involvement in controlling gene expression. This approach very clearly designated NELFE targets as enriched among the upregulated genes (Figure 2A). NELFE is a subunit of the Negative Elongation Factor (NELF) complex, which controls the speed and efficiency of RNAPII movement along the DNA, and regulates gene expression by causing RNAPII pausing downstream of the transcription start site (TSS). As all RNAPII-transcribed genes require NELF, annotated NELF-targets designate genes at which promoter escape rather than transcription initiation is the rate-limiting step (21).

**Figure 2:**
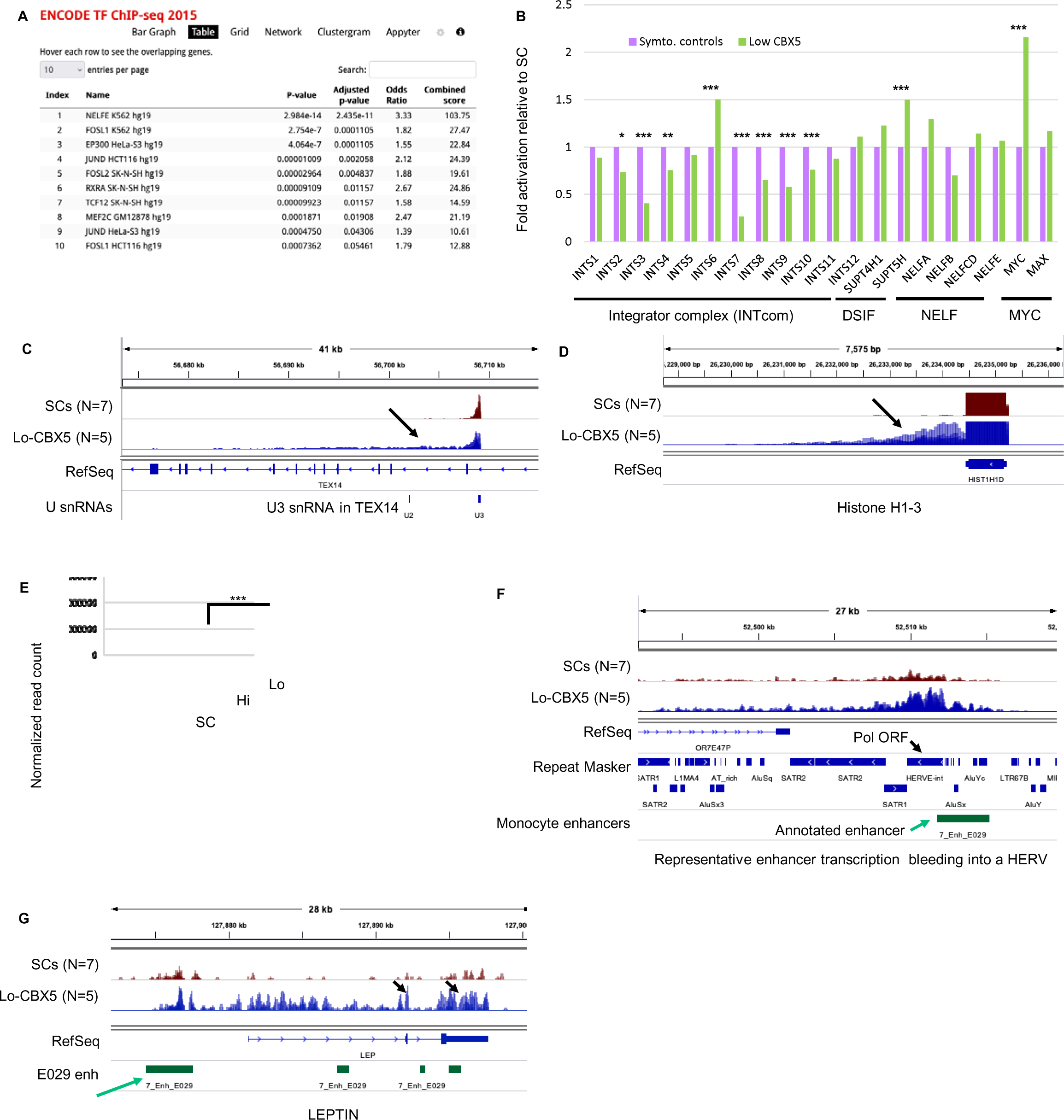
Evidence for a defective Integrator activity in Lo-CBX5 patients. (A) Analysis of genes down-regulated between SC and Lo-CBX5 patients using the ENCODE TF ChIP-seq data resource providing insights into the binding patterns of various transcription factors. (B) Bar graph reporting variation in expression of the indicated genes based on RNA-seq data in from SCs and Lo-Cbx5 patients. Levels in SCs set at 1. *: Adj. pVal<0.05, **: Adj. pVal<0.01, ***: Adj. pVal<0.001. (C-D) Screen shots from the IGV genome browser representing expression of the U3 snRNA copy in the TEX14 gene and of histone H1-3 (HIST1H1D) in SCs and Lo-CBX5 patients as indicated. (E) Quantification of reads mapping inside regions annotated as enhancers in monocytes by the Epigenome Roadmap. (F-G) Screen shots from the IGV genome browser representing expression of enhancers producing eRNAs either covering several repeated sequences, including a HERV (F), or resulting in transcription of the entire LEP gene (G).

The activity of NELF depends on the presence of at least two other complexes, the Integrator complex (INTcom) and the DRB Sensitivity-Inducing Factor (DSIF) complex (22). In addition, the negative impact of these complexes on transcription is balanced out by factors having a positive effect on elongation, including the Positive Transcription Elongation Factor b (P-TEF-b) and the transcription factor MYC (23).

Examination of the transcriptome did not indicate any clear variations in expression of the subunits of NELF. In contrast, 6 out 12 subunits of the INTcom were significantly (adj pVal<0.05) down-regulated (Figure 2B). Among these, INTS8 was previously reported to harbor single nucleotide polymorphisms (SNPs) associated with an elevated risk of MS (24). The INTcom was initially characterized for its function in the 3’-end processing of U snRNAs, during which it recognizes the 3’ cleavage site and facilitates the cleavage reaction (25). In that context, we noted that an earlier study had reported aberrant U snRNA polyadenylation in patients with MS, a known manifestation of a defective maturation of these RNA species (26). Examination of our patient RNA-seq data with a genome browser revealed increased accumulation of 3’ extensions at U snRNA transcription at multiple loci, characteristic of a hampered maturation (Figure 2C and Sup. Figure 2A). A compromised production of U snRNAs was also suggested by a defect in the maturation of several replication-dependent histone mRNAs, a process that relies on the U7 snRNA (27) (Figure 2D).

In parallel, the INTcom also promotes enhancer RNA (eRNA) instability, recognizing and cleaving their 3’-end (28). We therefore quantified RNA-seq reads within annotated monocyte enhancers, revealing a significant increase in eRNA accumulation in Lo-CBX5, but not in Hi-CBX5 patients (Figure 2E). In this analysis, we also quantified RNA-seq reads mapping to DNA repeats, and neither SINEs, LINEs, or endogenous retroviruses (HERVs) were upregulated genome-wide in the patients (neither Lo-CBX5s, nor Hi-CBX5s) (Sup. Figure 2B, 2C, 2D). The unaltered transcription of DNA repeats also documents that the low levels of CBX5 did not result in an overall restructuring of chromatin-mediated transcriptional silencing. We noted however an upregulation of HERVs coopted as enhancers, with transcription occasionally bleeding into DNA sequences encoding HERV-derived open reading frames (example in Figure 2F and quantification in Sup. Figure 2E). This suggested that the commonly observed upregulation of HERVs in patients with MS might be attributed to a deficiency in terminating or processing eRNAs that originate from viral regulatory regions coopted as enhancers. A similar phenomenon also seemed responsible for expression of protein coding genes. For example, we noted that the leptin (LEP) gene, normally detected only in adipose tissue and encoding a secreted hormone, was strongly activated in the Lo-Cbx5 patient monocytes (6-fold increased expression, Adj. pVal<0.03), apparently as a consequence of eRNAs originating from an upstream enhancer elongating into the LEP gene body, and undergoing splicing (Figure 2G). Thus, the modified turn-over of eRNAs appears as an unexpected source of ectopic gene expression in patients with MS.

### Modified RNAPII pause-release affects expression and splicing of genes relevant for MS

CBX5 activity is best documented in heterochromatin. However, CBX5 ChIP-seq data from HepG2 cells documented a clear enrichment of this chromatin-bound protein at transcription start sites (TSSs), suggesting that the chromatin regulator may also play a role in transcription initiation of protein-coding genes (Sup. Figure 3A). At these genes, INTcom participates in the regulation of RNAPII pause-release immediately downstream of the TSSs, and reduced activity of INTcom results in RNAPII entering the gene with reduced competence for elongation. This translates into exacerbated transcription at 5’ regions of genes, a phenomenon best observed with sequencing approaches detecting nascent pre-mRNA (29). However, visualization is also possible with RNA-seq data (30). To favor visualization of nascent RNAs in our RNA-seq patient data, we selected a set of genes with first introns exceeding 20kb in length, allowing the monitoring of reads originating from pre-mRNA over a long, uninterrupted region. Average transcription profiles over these genes revealed consistently increased accumulation of reads downstream of the TSS in the Lo-CBX5 patients as compared to SCs and Hi-CBX5 patients, thereby mirroring the pattern observed in earlier in vitro studies (Figure 3A and (29)). This accumulation was not seen at the 3’ end of genes (Figure 3B).

**Figure 3:**
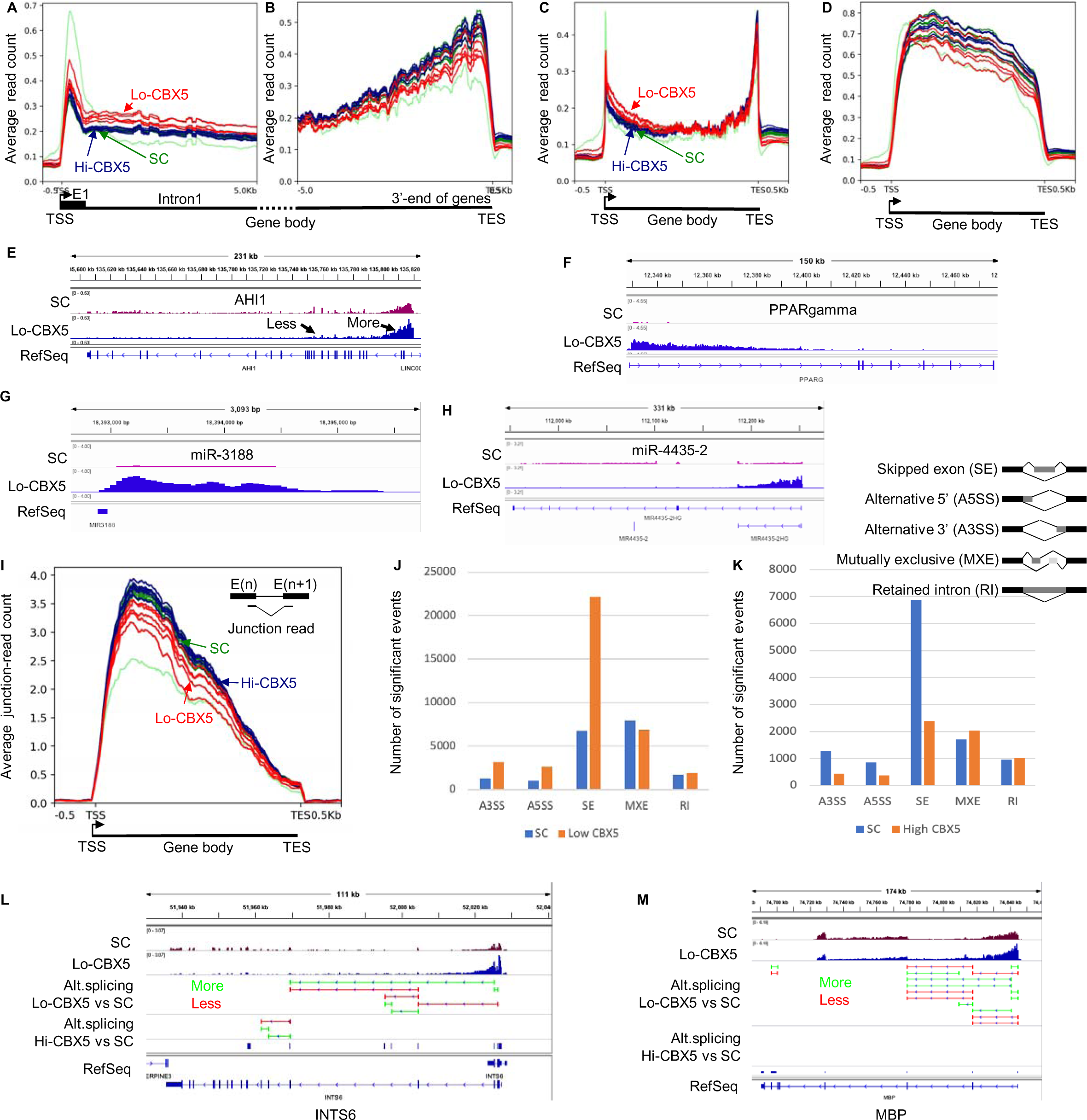
Defective Integrator activity results in extensive gene deregulation. (A-D) Average distribution of RNA-seq reads at genes containing an initial intron exceeding 20 kilobases in length. Profiles are either anchored on the Transcription Start Site (TSS in A), on the Transcription End Site (TES in B), or are plotted over the entire metagene either including (C), or excluding (D) intronic sequences. SC: symptomatic controls, Hi-CBX5: patients in the upper quartile for CBX5 expression levels, Lo-CBX5: patients in the lower quartile for CBX5 expression levels. (E-H) Screen shots from the IGV genome browser representing the impact of deregulated RNAPII pause-release at the indicated genes. (I) Average distribution of junction reads from the RNA-seq data across a metagene as in (C). (J-K) The RNA-seq was analyzed with the Rmats to quantify alternative splicing occurring between SC and Lo-CBX5 patients (J), or between SC and Hi-CBX5 patients (K). Histograms indicate number of significant events (Pval<0.05). A3SS: alternative 3’ splice site, A5SS: alternative 5’ splice site, SE: skipped exon, MXE: mutually exclusive exons, RI: retained introns. (L-M) Screen shots from the IGV genome browser representing at the indicated genes, splicing events either upregulated (green), or down-regulated (red) in the Lo-CBX5 (top track) or in the Hi-CBX5 (bottom track) when compared to SCs (Pval<0.05).

As an alternative method to visualize the phenomenon, we generated meta-genes with and without intronic sequences. When intronic sequences are excluded, the profiles show almost exclusively mature mRNA, more stable and therefore more abundant than pre-mRNA. Inversely, when including all sequences, the mature mRNAs account for only a minor fraction of the signal as exons are in average 300 nucleotides in length, while introns are several kilobases. These profiles confirmed increased transcription in the 5’ half of the genes in the Lo-CBX5 patients as compared to SCs and Hi-CBX5 patients (Figure 3C). In contrast, the mRNA output of the genes was in average equivalent in all donors (Figure 3D). Finally, we noted that genes activated in Lo-CBX5 patients were in average shorter than the down-regulated genes, a possible manifestation of globally increased transcription of regions located immediately downstream of the TSS (Sup. Figure 3B).

Consistent with these profiles, examination of the RNA-seq data with a genome browser allowed identifying multiple genes relevant for MS and displaying increased transcription over the first exon and the first intron, with consequences on their global level of expression. For example, the AHI1 gene, for which reduced expression has previously been associated with increased risk of MS (31), showed a clear increase in transcription down-stream of the TSS (“More” arrow, Figure 3E), while transcription was reduced from exon 5 and onwards (“Less” arrow, Figure 3E). This suggested that reduced control of RNAPII pause-release at this gene resulted in decreased production of full-length pre-mRNA. At other genes, uncontrolled promoter escape resulted in a net increase in the production of mature mRNA. This was exemplified by the PPARG and SLC7A11 genes, mostly transcribed in their 5’-regions, yet overall activated respectively 6-folds and 3-folds according to DESeq2 quantification (Adj. pVal<0.05 - see representative patients in Figure 3F and Sup. Figure 3C). The strong initiation/poor elongation behavior of the RNAPII also modified expression of regulatory RNAs when embedded in larger genes. For example, it caused increased expression of miR-3188 previously described as an MS risk factor (32,33) (Figure 3G), while resulting in decreased expression of miR-4435-2, a miR located further inside its host gene (Figure 2H).

From earlier studies, MS is known also to affect alternative splicing (AS) at many genes (34). Therefore, we examined whether the MS patients exhibited any alterations in overall splicing levels, using the RNA-seq data to quantify junction reads (reads produced when the sequenced RNA fragments span over exon-intron boundaries - such reads serve as evidence of a splicing event). This approach showed that, on average, less splicing events were detected in Lo-CBX5 patients when compared to Hi-CBX5 and SC donors, with a more pronounced impact at the 5’-end of genes (profiles Figure 3I, red below green and blue). Further analysis of the RNA-seq data with the RMATs package confirmed that alternative splicing was affected in the MS patients, whether displaying Lo- or Hi-CBX5 expression. Yet, it also documented that the reduced splicing level in the Lo-CBX5 patients essentially translated into exon skipping, consistent with the fact that fewer splicing events are needed when fewer exons are included (Figure3J). Inversely, Hi-Cbx5 patients displayed reduced exon skipping when compared to the SCs (Figure 3K). Interestingly, increased exon skipping in Lo-CBX5 patients was observed at the INTS6 gene, suggesting that reduced Integrator activity is a self-reinforcing phenomenon (Figure 3L). Increased exon skipping was also observed at MBP and GAPDH genes encoding MS auto-antigens (Figure 3M, and Sup. Figure 3D).

Thus, reduced Integrator activity translates into either up- or down-regulation of numerous genes associated with multiple sclerosis (MS), the outcome possibly varying depending on the extent to which these genes rely on RNAPII pause-release for their regulation. In addition, gene expression is further impacted by overall reduced splicing.

### Inactivation of Cbx5 in the mouse promotes inflammation and exacerbates EAE

To explore further a possible causative role for reduced CBX5 activity in the MS pathogenesis, we implemented a mouse model inactivated for the Cbx5 gene. This mouse model was viable, although animals homozygote for the Cbx5 mutation (Cbx5-/-) displayed high perinatal mortality.

We first challenged the mouse model with Experimental Autoimmune Encephalomyelitis (EAE), a protocol widely used to mimic some aspects of MS by triggering an autoimmune response with injections of a myelin-derived peptides followed by an induction of inflammation by administration of pertussis toxin (Figure 4A). Cbx5-/- mice displayed an exacerbated reaction to that protocol, with an earlier onset of the symptoms followed by a rapid reaching of high grade EAE, beyond recovery (Figure 4B). The early onset and the exacerbated symptoms were also observed with heterozygote Cbx5+/- mice and these animals experienced a more extensive loss of body weight than the wild-types during the protocol (Sup. Figure 4A and 4B). Unlike the Cbx5-/-, the Cbx5+/- mice eventually transitioned into a recovery phase, although with a one-day delay compared to the wild-types (Sup Figure 4A).

**Figure 4:**
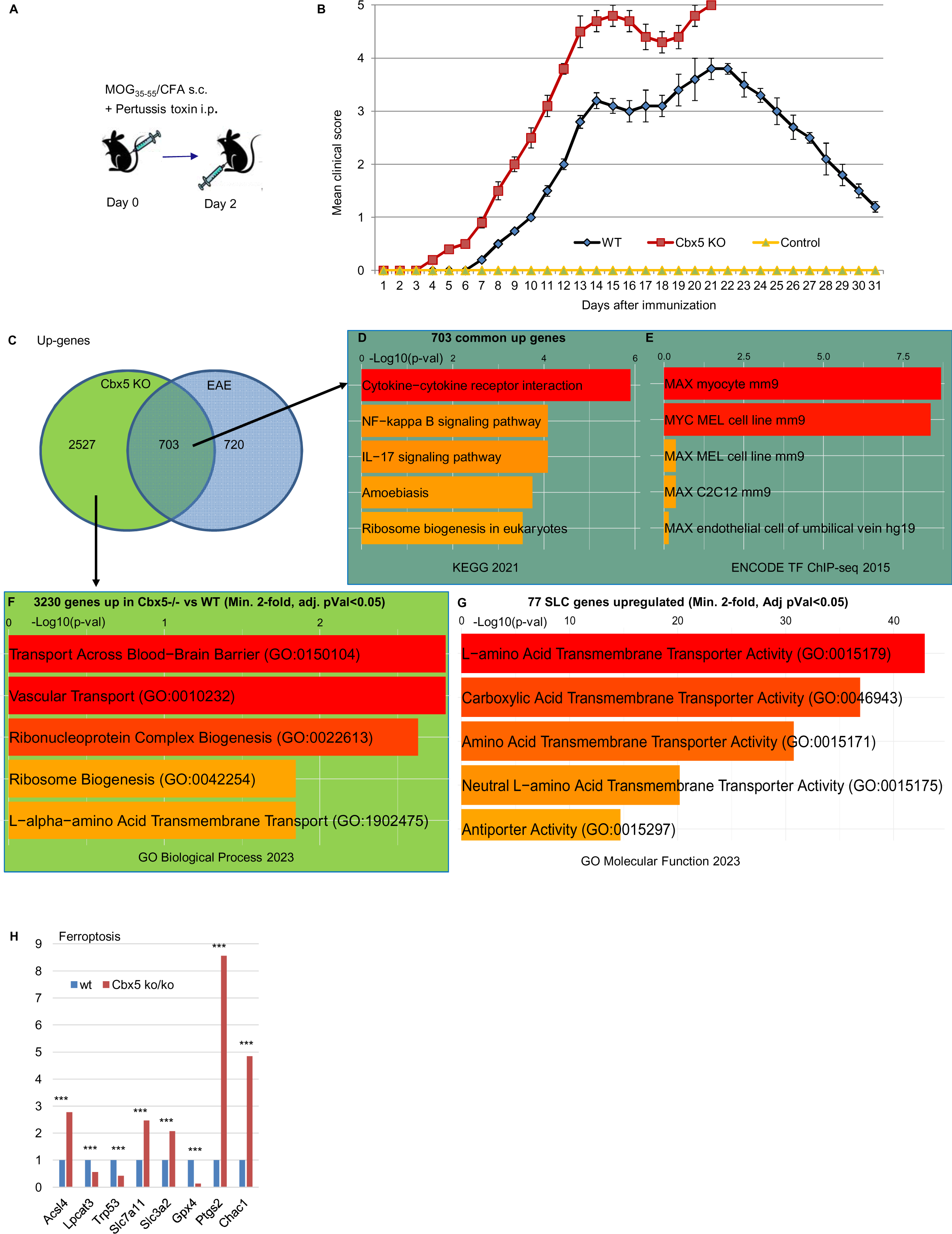
Inactivation of Cbx5 in the mouse exacerbates EAE. (A) Schematic representation of the EAE protocol. (B) Graphic representation of the clinical scores observed for the mice included in the EAE protocol. 0: No clinical symptoms. 1: Flaccid tail (loss of tail tone). 2: Hind limb weakness or partial paralysis. 3: Complete hind limb paralysis. 4: Complete hind limb paralysis and partial forelimb paralysis. 5: Moribund or dead. Indicated data are averaged from 5 females, 10 weeks of age. (C) Venn diagram representing the number of genes up-regulated either in Cbx5 null mice compared to WT, in mice exposed to an EAE protocol compared to controls, or in both conditions. (D-E) Analysis of genes up-regulated both upon Cbx5 inactivation and by the EAE protocol using the KEGG database providing information about the molecular functions and biological pathways of genes (D), and the ENCODE TF ChIP-seq data resource providing insights into the binding patterns of various transcription factors (E). (F) Analysis of genes up-regulated upon Cbx5 inactivation using the "Biological Process" category in the GO database focusing on describing biological events, pathways, and processes. (G) Analysis of the 77 SLC genes up-regulated (Min. 2-fold, Adj pVal<0.05) upon Cbx5 inactivation using the "Molecular Function" category in the GO database focusing the molecular activities of gene products. (H) Bar graph reporting variation in expression of the indicated genes based on RNA-seq data in from WT and Cbx5-/- mice. Levels in WTs set at 1. *: Adj. pVal<0.05, **: Adj. pVal<0.01, ***: Adj. pVal<0.001.

We next carried out RNA-seq on CD4+ T cells from either WT (n=2), Cbx5+/- (n=1), Cbx5-/- (n=3), EAE (n=3), and Cbx5-/- EAE (n=3). As anticipated from the Lo-CBX5 MS patients, inactivation of Cbx5 had an extensive impact on the transcriptome, with a total of 6478 differentially regulated genes (3224 up and 3248 down genes 2-fold or more, Adj pVal<0.05 – Sup. Figure 4C and 4D and Sup. Tables 2A, 2B, and 2C). Among the 3224 upregulated genes, 703 genes were also upregulated by EAE, corresponding to approximately half of the genes upregulated by EAE (Figure 4C). This observed intersection was approximately 6-fold higher than expected by chance (Sup Figure 4E). Pathway analysis on the 703 shared genes revealed that Cbx5-/- mice activated IL-17 and NFkappaB pathways at steady state, which may participate in the exacerbated response to the EAE protocol (Figure 4D). We noted also that the 703 genes upregulated both by Cbx5 inactivation and by EAE were enriched in Myc target genes (Figure 4E). This observation suggested a potential reliance of these genes on regulation through RNAPII pause-release mechanisms (35).

Gene ontology analysis of the 3224 genes upregulated following Cbx5 inactivation identified strong enrichment scores for transmembrane transport through the blood-brain barrier, underscoring the modified expression of numerous solute carrier (SLC) genes (Figure 4F). These genes encode membrane transport proteins involved in the movement of ions, nutrients, and metabolites and frequently associated with inflammation and autoimmunity disease (36). We noted that among the 373 SLC genes annotated in the mouse genome, 117 displayed a modified expression (min. 2-fold, Adj. pVal<0.05), while the 77 upregulated SLC genes were enriched in amino acid transporters (Figure 4G). These observations suggest that SLC genes are particularly sensitive to Cbx5 regulation, and that the modified expression of the SLC genes may promote T cell activation, heavily relying on amino acid transportation (37).

To mirror our study on the MS patients, we also examined the ferroptosis pathway. We here noted a strong down-regulation of Gpx4 (10-fold) and a corresponding upregulation of the ferroptosis markers Ptgs2 and Chac1 (Figure 4H). Consistent with a possibly increased cell death by ferroptosis, we noted also that down-regulated genes (down 2-fold or more, Adj pVal<0.05, 3248 genes) were enriched in genes involved in fatty acid beta oxidation (FAO – Sup. Figure 4F). A down-regulation of this pathway is anticipated to favor ferroptosis, as FAO normally consumes the fatty acids, leading to a reduction in the rate of lipid peroxidation (38). Analysis of the down-regulated genes with either Reactome, or BioPlanet databases revealed that these genes were also enriched in genes involved in cell cycle regulation, DNA replication, and mitochondrial ATP synthesis, consistent with a subset of the cells possibly withdrawing from the cell cycle (Sup. Figure 4G and 4H).

### The Cbx5-/- mouse model recapitulates the defect in Integrator activity

Finally, we investigated whether Cbx5-/- mice recapitulated some of the transcriptional anomalies observed in Lo-CBX5 MS patients. Examination of the transcriptome revealed that, as observed for the Lo-CBX5 patients, regulators of snRNA/eRNA processing, and RNAPII pause-release were affected, with reduced expression of several INTcom subunits (Ints3, Ints4, Ints9, Ints10, and Ints11/Cpsf3l), and of the DSIF subunit Supt4h1/Spt4, while expression of Myc was increased (Figure 5A). The RNA-seq data also suggested that the functions of these regulators were affected. At U snRNA and histone genes, we noted that despite a decrease in overall expression, the read-count in the 3’ regions was relatively higher, suggestive of a reduced 3’ end processing (see examples Figure 5B and Sup. Figure 5A). To probe for defects in the regulation of RNAPII pause-release, we plotted the average distribution of reads at a set of genes harboring a first intron of more than 20kb, thereby using the same approach as for the MS patient samples. For the 3 Cbx5-/- samples and the one Cbx5+/-, alike what we observed in the Lo-CBX5 patients, accumulation of reads was increased over the region downstream of the TSS, while unaffected or reduced over 3’ region of the genes (Figures 5C and 5D). The increased transcription at the TSS-proximal regions was also visualized when plotting the distribution over an average (meta) gene including both exons and introns (Figure 5E). In contrast, plotting the read distribution over exons only, revealed an overall reduced production of mature mRNA, consistent with the mouse model recapitulating a reduced efficiency of splicing (Figure 5F). The Ints6 gene provided a clear illustration of this transcriptional defect, with accumulation of intronic reads, associated with a reduced signal at exons, particularly toward the 3’ end of the gene (Figure 5G). Analysis of the data with the RMATS package confirmed an extensive impact of Cbx5 inactivation on alternative splicing, with, as observed in the Lo-CBX5 patients, an increase in exon-skipping events. Yet, consistent with an accumulation of non-maturated pre-mRNA species, the most favored type of alternative splicing was intron retention (Figure 5H). EAE alone reduced rather than increased accumulation of reads on the 5’ region of genes (Sup. Figure 5B). Yet, like in the untreated animal, Cbx5 inactivation resulted accumulation of reads downstream of the TSSs (Sup. Figure 5C).

**Figure 5:**
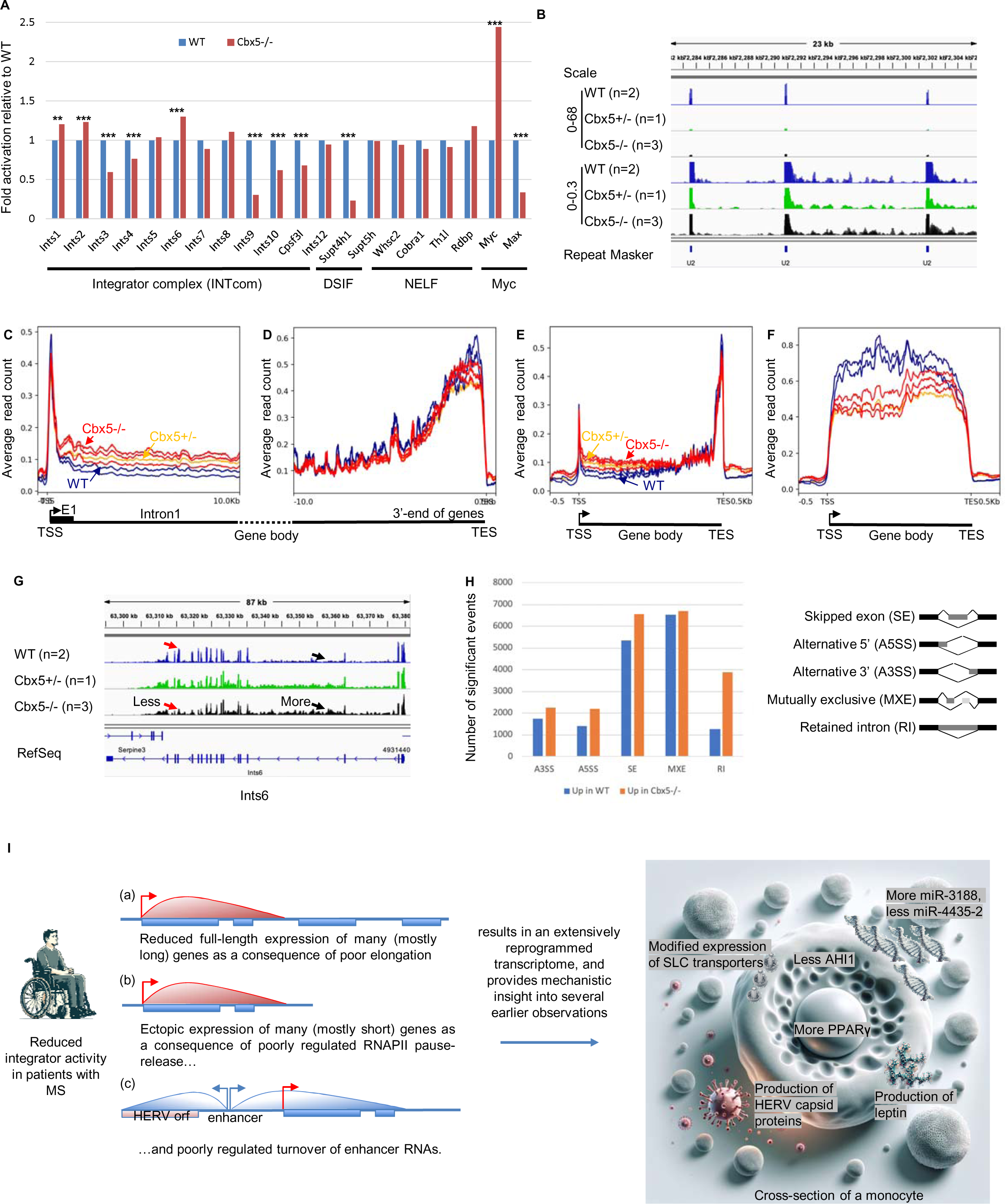
The mouse model confirms the impact of Cbx5 on Integrator activity. (A) Bar graph reporting variation in expression of the indicated genes based on RNA-seq data in from WT and Cbx5-/- mice. **: Adj. pVal<0.01, ***: Adj. pVal<0.001. (B) Screen shots from the IGV genome browser representing expression of at a series of U2 snRNA copies in WT, Cbx5+/-, and Cbx5-/- mice as indicated. Schematic shows two different scales to allow visualization of the reduced expression of the U2 snRNAs (top tracks) and of the downstream reads accumulating at similar levels in all samples (bottom tracks). (C-F) Average distribution of RNA-seq reads at genes containing an initial intron exceeding 20 kilobases in length. Profiles are either anchored on the Transcription Start Site (TSS in C), on the Transcription End Site (TES in D), or are plotted over the entire metagene either including (E), or excluding (F) intronic sequences. (G) Screen shots from the IGV genome browser representing the impact of deregulated RNAPII pause-release at the Ints6 gene. (H) Histograms indicate number of significant events (Pval<0.05). A3SS: alternative 3’ splice site, A5SS: alternative 5’ splice site, SE: skipped exon, MXE: mutually exclusive exons, RI: retained introns. (I) Model: A subset of patients with MS, particularly those in progressive phases of the disease, exhibit reduced Integrator activity. This leads to a loss of control over RNAPII pause-release, resulting in increased transcription at promoter-proximal regions and decreased transcription in more distal regions. Consequently, this causes reduced expression of some genes (a) and activation of others (b), with the impact depending not only on the size of the gene but also on the reliance of each gene on regulation at the level of RNAPII pause-release. In parallel, the reduced Integrator activity also affects the control of eRNA maturation (c), and transcripts initiated at enhancers occasionally extend into neighboring genes, eventually of viral origin (GAG, Pol, Env). These transcriptional anomalies may explain numerous transcriptional events previously associated with MS, including the production of retrovirus-like particles, altered leptin production, changes in the expression of various miRNAs, altered expression of SLC11A7/xCT and AHI1, and increased expression of PPARγ.

## DISCUSSION

Our RNA-seq analysis of a relatively diverse patient cohort revealed that CBX5 expression levels, rather than comorbidity, age, or gender, partitioned patients into two distinct groups. Patients with low CBX5 expression showed significant transcriptional differences from control donors, whereas those with high CBX5 expression resembled the controls more closely, with a transcriptome mainly characterized by reduced expression of vitamin D-regulated genes. The correlation between the level of serum vitamin D and disease activity still needs to be further investigated (39). Yet, this observation suggests that the Hi-CBX5 group, which included individuals with clinically and radiologically isolated syndromes, might particularly benefit from a vitamin D-enriched diet.

The Lo-CBX5 group was enriched in patients in primary and secondary progressive stages, while devoid of isolated syndromes, suggesting that low CBX5 expression may be indicative of a steady neurological decline. At the transcriptional level, the group was characterized by defects in U snRNA and enhancer-RNA (eRNA) processing, as well as in RNA polymerase II (RNAPII) pause-release, collectively indicative of reduced Integrator complex activity. An implication of the Integrator complex in the MS pathology has hitherto not been investigated, although several indications have been pointing in that direction. Firstly, single nucleotide polymorphisms (SNPs) within the Integrator subunit INTS8 have previously been associated with an elevated risk of MS (24). Another study has also detected association between MS and SNPs in genes associated with “abortive elongation”, including the NELF subunits NELFE and NELFCD, and the DSIF subunit SUPT4H1 (40). Finally, patients with MS have been reported to show aberrant U snRNA polyadenylation, a known manifestation of a defective maturation of these RNA species (26), while we have previously described increased accumulation of eRNAs associated with MS (11). More generally, we note that the integrator complex is a known player in neurological diseases, with mutations in INTS1 and INTS8 being associated with rare recessive human neurodevelopmental syndromes (41).

Reduced Integrator activity affects expression of hundreds of protein-coding genes independently of its impact on snRNA maturation (29,42). By restricting pause-release to elongation-competent RNAPII enzymes, the Integrator complex promotes production of full-length pre-mRNAs. Conversely, limited Integrator activity biases transcription towards promoter-proximal regions (illustrated in model Figure 5I). A distribution of reads indicative of diminished Integrator activity was observed at the AHI1 gene, peaking at the initial exons, then rapidly decreasing (Figure 2E). This suggests that the reduced expression of AHI1, which is strongly linked to a higher risk of MS, might be due either to SNPs as previously reported (31), or to decreased Integrator activity. In parallel, poorly controlled RNAPII pausing upregulates basal transcription of (often short) inducible genes (43). Therefore, the chronic activation of short intron-less pro-inflammatory genes such as c-Jun and JunD may be directly related to this mechanism, consistent with up-regulated genes being in average shorter than down-regulated genes in the Lo-CBX5 patients. Longer genes affected by this mechanism include PPARG and SLC7A11 (xCT) and may be activated in the context of a protective effect, PPARG modulating the immune response (44), while SLC7A11 promoting cell death as will be discussed below (45). Finally, the RNAPII incompetent for elongation also accounts for the observed downregulation of miR-3188 and upregulation of mir-4435-2, respectively located either at the start of the host gene, receiving excessive RNAPII, or located in a more central regions, thus being out of reach of the polymerase (32,33).

Deregulated elongation of enhancer RNAs (eRNAs), another hallmark of reduced Integrator activity, emerges as a second mechanism promoting ectopic gene expression (refer to model in Figure 5I). A prime example of this is the increased transcription of HERV-encoded sequences. Promoter regions of ancient viruses inserted in the human genome are occasionally coopted as enhancers (46). We observe that at some of these enhancers of viral origin, the increased elongation of eRNAs mechanically results in the transcription of the adjacent viral genes. This phenomenon may possibly explain the increased expression of HERV envelope proteins at the surface of monocytes and the production of retrovirus-like particles from cultured patient cells (47,48). The impact of elongated eRNAs was also exemplified by the leptin gene that we found transcribed from an upstream enhancer bleeding into the LEP gene. This adipokine normally secreted by adipose tissues regulates systemic metabolism and appetite, while also signaling directly to immune cells to promote inflammation (49). Leptin has been found to be present at increased levels in patients with MS, and has been associated with MS risk (50,51). This may directly illustrate how defective Integrator activity can participate in the pathogenesis of MS.

Reduced Integrator activity was the main transcriptional phenotype of the mouse T cells inactivated for Cbx5, suggesting that the encoded HP1α protein is directly involved in the regulation of RNAPII activity. This is in contrasts with previously documented heterochromatic activity of this protein. Yet, examination of the available ChIP-seq data revealed a localization of HP1α at the TSSs of genes. We note also that HP1α is known to dimerize with TRIM28/KAP1 previously reported to participate in the regulation of RNAPII elongation (52,53). Finally, HP1 proteins copurifies with the CBC cap binding complex, consistent with a role in early phases of transcription initiation (54). In an earlier study, we had shown that HP1α activity was affected by increased citrullination of histone H3R8 (10). In the monocytes of the cohort of MS patients examined here, we did not observe any increased transcription of PADI enzymes responsible for this citrullination event. This suggests that HP1α activity may be compromised via either transcriptional or post-transcriptional mechanisms, possibly as a function of the cell type under scrutiny.

The mechanism causing Cbx5-inactivation to promote EAE still needs to be explored at an immunological level to identify the involved cell populations. Yet, we note that the many genes up-regulated both by EAE and Cbx5 inactivation were enriched in Myc targets. The Myc transcription factor is indispensable for T cell activation, playing a role in the amino acid supply required for protein production associated with the increased cellular activity (55). The amino acid transporter Slc7a5, a key target of Myc, is also among the numerous SLC genes up-regulated by Cbx5 inactivation (> 4-fold, Adj. Pval=10^-80^). This apparently exacerbated sensitivity of genes of the SLC family to deregulated RNAPII pause-release may possibly explain some aspects of the increased responsiveness of Cbx5 null mice to EAE.

While displaying numerous similarities, the transcriptional defects observed in Lo-CBX5 patients and in Cbx5 null mice were not entirely identical. In particular, in the mouse model, we noted a reduced expression of U snRNAs and a more pronounced disruption of pre-mRNA maturation. It is plausible that these two phenomena are related. The significant scarcity of U snRNAs in the mouse model could be a major factor contributing to the extensive intron retention. In contrast, in patients, where the CBX5 deficit is milder, a relatively less affected splicing machinery may cause only increased exon skipping.

Whether reduced CBX5 activity is a trigger of MS symptoms remains an open question. We noted that in the Cbx5 KO mouse model, inactivation of just one Cbx5 allele was sufficient to obtain both the reduced Integrator activity and the exacerbated reaction to EAE. Nevertheless, the mice did not manifest EAE symptoms spontaneously, suggesting that Cbx5 inactivation generates a conducive terrain, but that the neuroinflammatory symptoms necessitate an external trigger to develop. Although our data do not offer insights into the potential nature of this trigger, they may provide valuable information regarding subsequent transcriptional events. Indeed, several observations suggest that reduced Integrator activity is a self-amplifying mechanism. First, mutations in INTS1 and INTS8 were previously shown to result in the down-regulation of multiple subunits of the INTcom (41). A similar phenomenon was observed upon Cbx5 inactivation in our mouse model, with down-regulation of Ints3, Ints4, Ints9, Ints10, and Cpsf3l/Ints11. The reduced expression of these INTcom genes may be due to a sensitivity of these genes to their own activity, as we documented for the INTS6 gene in both human and mouse. This suggests that external stimuli hitting on RNAPII pause-release may trigger a feed-forward loop resulting in reduced INTcom activity. In this context, we note that the Epstein Barr virus (EBV) was found associated to Multiple Sclerosis (MS), first described 30 years ago (56) and has continued to show evidence as a causal role in MS (57). This virus is known to target INTS6 (58), while also being a promoter of ferroptosis (59). Together with the clear evidence for ferroptosis in the Cbx5-/- mice, this may suggest that ferroptosis is the normal fate of lymphocytes experiencing a defective Integrator activity. The seemingly abortive ferroptosis in the MS patients may further suggest that this defense mechanism is not entirely functional in these patients, possibly participating in the exacerbated neuroinflammation.

## MATERIAL AND METHODS

### Ethics statement

The study was conducted in accordance with the Ethical Declaration of Helsinki and all patients gave written, informed consent. The study and the material for informed consent were approved by The Central Denmark Region Committee on Biomedical Research Ethics (journal number: 1-10-72-334-15).

### Patients and controls

Patients admitted to the MS clinic, Department of Neurology, Aarhus University Hospital, were consecutively included from January 2017 to January 2018. A full diagnostic workup included medical history, clinical examination, MRI of the entire neural axis, cerebrospinal fluid (CSF) analyses (cells, protein, IgG index, oligoclonal bands) and evoked potentials as recommended (60). CSF and MRI examination were evaluated according to the revised MacDonald criteria from 2017 (60) and an Extended Disability Status Scale score was assessed according to Kurtzke (61). Patients were excluded if they had other neurologic disease or received glucocorticoids within the month preceding sampling. Total number of MRI white matter lesions were registered by fluid-attenuated inversion recovery sequences on MRI. Demographics and paraclinical findings of patients with CIS, RRMS, PPMS, RIS and SCs are summarized in Table 1. Patients included as SC have neurological symptoms, but have no objective clinical or paraclinical findings to define a specific neurological disease. This specific definition is described in detail by Teunissen et al (62) and they do not represent early MS.

Patients included as RIS have no neurological symptoms and are only referred to diagnostic workup owing to the presence of incidental white matter lesions in MRI suggestive of MS. Diagnostic criteria for RIS were proposed in 2009 and include the number, shape and location of the brain lesions (60). Lesions are ovoid and well circumscribed with a size greater than 3 mm, show dissemination in space and can be juxtaposed to the corpus callosum. Lesions should not follow a vascular distribution and do not account for any other pathologic processes.

### Data download

We downloaded the Cbx5 ChIP-seq data acquired in hepatocarcinoma-derived HepG2 cells from the ENCODE portal, output type was “signal p-value” in bigwig format, file ENCFF408KOE (https://www.encodeproject.org/)(63).

### RNA sequencing

Monocytes were obtained from patient PBMCs using a Pan Monocyte Isolation Kit (Miltenyi Biotech, ref. 130-096-537) following the kit protocol, allowing for negative selection of unstimulated monocytes. CD4+ mouse T cells were purified from spleenocytes using the Miltenyi kit 130-104-454. For both cell types, total RNA was extracted by Trizol LS (Thermo Fisher Scientific, ref. 10296028), according to the manufacturer’s protocol. Total RNA library preparation and sequencing were performed by Novogene Co., Ltd, as a lncRNA sequencing service, including lncRNA directional library preparation with rRNA depletion (Ribo-Zero Magnetic Kit), quantitation, pooling and PE 150 sequencing (30G raw data) on Illumina HiSeq 2500 platform. Filtering and trimming of the RNA seq data left around 230-300 million reads pairs/sample. Mapping was carried out with STAR (v2.6.0b) (parameters: --outFilterMismatchNmax 1 --outSAMmultNmax 1 --outMultimapperOrder Random -- outFilterMultimapNmax 30) (64). The reference genomes were hg19 homo sapiens and mm10 mus musculus primary assemblies from Ensembl. The SAM files were converted to BAM files and sorted by coordinate using samtools (v1.7) (65).

### BigWig files, heatmaps, and profiles

Bigwigs files were generated from .bam files with bamCoverage (parameter: -- normalizeUsing CPM) from Deeptools (v3.1.3) (66). Heatmaps and profiles were also generated with Deeptools (v3.1.3). Matrices were generated with computeMatrix followed by plotProfile or plotHeatmap as appropriate.

### Data quantification

Read quantification was carried out with featureCounts (v1.6.1) from the Subread suite (67). For repeated element, files were obtained by extracting in .bed file format, entries annotated “SINE”, “LINE” or “LTR” in the “RepClass” field from RepeatMasker.

### Analysis of splicing

Differential splicing analysis was done using rMATS (v4.1.0) (68) with the parameters: -- libType fr-firststrand –novelSS. The occurrence of each type of splicing event was then counted.

### Pathway analysis and data visualization

Pathway analysis was carried out with Enrichr from the Ma’ayan lab (69,70). The Integrative Genomics Viewer software (IGV) was used to examine specific loci (71).

### Cbx5 null mouse model

C57BL/6N-Atm1Brd Cbx5tm1a(EUCOMM)Wtsi/WtsiOrl strain inactivated for Cbx5 was received from the EUCOMM Consortium in the context of a Standard Material Transfer Agreement. It was transferred via the TAAM – CNRS Orléans for breeding at the Institut Pasteur - Paris. The experiments involving the mouse model were carried out in strict accordance with the approved research protocols and in compliance with all applicable institutional and regulatory guidelines (CETEA Institut Pasteur approval dap170030).

### Experimental Autoimmune Encephalomyelitis (EAE)

EAE was carried out as previously described (72). Briefly, to prepare the MOG35-55 emulsion for immunization, we first calculated the total volume required and then prepared 1.5 to 2 times that amount to account for potential losses during the process. Each mouse received a subcutaneous injection of 200 µl of a mixture containing MOG35-55 peptide solution and Complete Freund’s Adjuvant (CFA) in a 1:1 ratio. The MOG35-55 peptide solution was prepared by diluting lyophilized 200 µg of the peptide per mouse with ddH2O to achieve a final concentration of 2 mg/mL, which was then stored at -20°C. CFA was prepared by grinding 100 mg of desiccated Mycobacterium tuberculosis H37RA and adding 10 mL of Incomplete Freund’s Adjuvant (IFA) to create a 10 mg/mL CFA stock solution that could be stored at 4°C. Just prior to immunization, CFA was diluted with IFA to reach a final concentration of 2 mg/mL, with thorough mixing. The MOG35-55 and CFA were then mixed in a 1:1 ratio until the final concentration of 1 mg/mL was achieved. This emulsion, a critical step for immunization, was carefully prepared to ensure proper emulsification. After verification of its stability, the emulsion was drawn into syringes for subsequent use. In addition to the MOG35-55 emulsion, pertussis toxin was prepared and administered. For each mouse, 400 ng of pertussis toxin in 200 µl of PBS was injected intraperitoneally on the day of immunization and repeated two days later. Pertussis toxin was reconstituted by dissolving 50 µg in 500 µl of ddH2O to create a 100 µg/mL stock solution, which was stored at 4°C. To achieve the required concentration for immunization, the stock solution was diluted 1:50 with PBS. Finally, to ensure proper identification and monitoring, individual mice were marked, typically by using color markings on the tail base. A second dose of pertussis toxin was administered on day two after the initial immunization to complete the immunization process.

### Blood brain barrier permeability

In the conducted experiments, mice were administered Fluorescein-cadaverine via tail vein injections at a dosage of 6 μg/g. Following a 2-hour waiting period, the mice were anesthetized, and their brains were perfused with ice-cold PBS for 7 minutes. The cortex was subsequently isolated and weighed, and cortical tissues were homogenized using a 1% Triton X-100 in PBS solution. After centrifugation at 12,000 × g for 20 minutes at 4 °C, the resulting supernatant was used to measure fluorescence with specific excitation and emission wavelengths. Importantly, to ensure the accuracy of the fluorescence measurements, the obtained values were normalized by dividing them by the respective cortical weights. This meticulous methodology allowed for precise and standardized assessment of tracer leakage and fluorescence in these experiments, contributing to the reliability of the results.

### Data availability

All data in the figures are available in the published article and in online supplemental material. Human monocyte and mouse Cbx5 KO RNA-seq data are available at the Gene Expression Omnibus respectively under accession no. GSE249613 and no. GSE249605.

## ACKNOWLEDGMENTS

The work was supported by a grant from REVIVE, an ANR "Laboratoire d’Excellence" program (2011 - 2021). TC received funding from The Graduate School of Health, Aarhus University, The Danish MS Society, The Beckett Foundation, and Dagmar Marshalls Foundation. Travel expenses of M. Carstensen for the collaboration with CM were covered by The Knud Højgaards Foundation, The Augustinus Foundation, The Aage og Johanne Louis-Hansens Foundation, and The Oticon Foundation.

**Sup. Figure 1: Extensive transcriptional deregulation in monocytes from patients with reduced expression of CBX5** (A) Analysis of genes up-regulated in Lo-CBX5 patients when compared to symptomatic controls using the KEGG database providing information about the molecular functions and biological pathways of genes. (B) Bar graph reporting variation in expression of the indicated genes based on RNA-seq data in from SCs and Lo-Cbx5 patients. Levels in SCs set at 1. *: Adj. pVal<0.05, **: Adj. pVal<0.01, ***: Adj. pVal<0.001. (C) Analysis of genes down-regulated in Lo-CBX5 patients when compared to symptomatic controls using the BioPlanet or WikiPathway databases as indicated.

**Sup. Figure 2: Evidence for a defective Integrator activity in Lo-CBX5 patients** (A) Screen shots from the IGV genome browser representing expression of the U2 snRNA copy upstream of the WDR74 gene. (B-E) Quantification of reads mapping inside regions annotated as SINEs, LINEs, or LTRs in ReapeatMasker (B-D), or both as LTRs in ReapeatMasker and as enhancers in monocytes by the Epigenome Roadmap (E).

**Sup. Figure 3: Defective Integrator activity results in extensive gene deregulation** (A) Heatmap of HP1a/CBX5 ChIP-seq signal in HepG2 cells (SRR14105564). The heatmap is anchored on the Transcription Start Site of genes with an exon 1 longer than 20kb from the Hg19 version of the genome. (B) Bar graph representing the average size in nucleotides of the genes either upregulated (1395 genes), or down-regulated (1350 genes) in Lo-CBX5 patients compared to SCs. (C) Screen shots from the IGV genome browser representing the impact of deregulated RNAPII pause-release at the SLC7A11 gene. (D) Screen shots from the IGV genome browser representing at the GAPDH gene, splicing events either upregulated (green), or down-regulated (red) in the Lo-CBX5 (top track) or in the Hi-CBX5 (bottom track) when compared to SCs (Pval<0.05).

**Sup. Figure 4: Inactivation of Cbx5 in the mouse exacerbates EAE.** (A) Graphic representation of the clinical scores observed for the mice included in the EAE protocol either WT or Cbx5+/-. 0: No clinical symptoms. 1: Flaccid tail (loss of tail tone). 2: Hind limb weakness or partial paralysis. 3: Complete hind limb paralysis. 4: Complete hind limb paralysis and partial forelimb paralysis. 5: Moribund or dead. Indicated data are averaged from 10 females, 10 weeks of age. (B) Graphic representation of the body weight of either WT or Cbx5+/- mice included in the EAE experiment. Indicated data are averaged from 10 females, 10 weeks of age. (C-D) Vulcano plots illustrating differential gene expression between Cbx5-/- and WT mice either at steady state (C), or involved in an EAE protocol (D). (E) Intersection between genes upregulated upon Cbx5 inactivation and the 1423 genes upregulated by EAE was compared to the intersection between genes upregulated upon Cbx5 inactivation and a list of 1423 randomly selected genes (average of 10 iterations). (F) Analysis of genes down-regulated in Cbx5-/- mice compared to WT using the Go Biological Process, Reactome, or BioPlanet databases as indicated.

**Sup. Figure 5: The mouse model confirms the impact of Cbx5 on Integrator activity** (A) Screen shots from the IGV genome browser representing expression of histone gene H2ac25 in WT, Cbx5+/-, and Cbx5-/- mice as indicated. Schematic shows two different scales to allow visualization of the reduced expression of the histone (top tracks), and of the downstream reads accumulating at similar levels in all samples (bottom tracks). (B-C) Average distribution of RNA-seq reads at genes containing an initial intron exceeding 20 kilobases in length. Profiles are plotted over the entire metagene and compare either WT and WT EAE (B), or Cbx5-/- and Cbx5-/- EAE (C).

## Notes

### Competing Interest Statement

The authors have declared no competing interest.

### Summary of Updates

The quality of the figures has been improved and there has been some rewriting of the text.

