## Supplemental Figures for "Defective Integrator activity shapes the transcriptome of patients with multiple sclerosis"

**A** Genes **up** in Low CBX5 compared to **SC**: 1395 genes

**KEGG 2021 Human** Bar Graph **Table** Clustergram Appyter

Hover each row to see the overlapping genes.

10 entries per page Search:

| Index | Name | P-value | Adjusted p-value | Odds Ratio | Combined score |
|---|---|---|---|---|---|
| 1 | Rheumatoid arthritis | 0.0002152 | 0.02023 | 3.01 | 25.40 |
| 2 | HIF-1 signaling pathway | 0.0001802 | 0.02023 | 2.84 | 24.49 |
| 3 | MAPK signaling pathway | 0.0001743 | 0.02023 | 2.01 | 17.37 |
| 4 | Ferroptosis | 0.0004017 | 0.02832 | 4.33 | 33.83 |
| 5 | IL-17 signaling pathway | 0.0007432 | 0.03342 | 2.76 | 19.86 |
| 6 | TGF-beta signaling pathway | 0.0007432 | 0.03342 | 2.76 | 19.86 |
| 7 | NF-kappa B signaling pathway | 0.0008295 | 0.03342 | 2.63 | 18.63 |
| 8 | Legionellosis | 0.001715 | 0.04836 | 3.21 | 20.42 |
| 9 | AGE-RAGE signaling pathway in diabetic complications | 0.001471 | 0.04836 | 2.56 | 16.69 |
| 10 | Fluid shear stress and atherosclerosis | 0.001574 | 0.04836 | 2.26 | 14.58 |

**B** Ferroptosis pathway genes in **Low** CBX5 compared to **SC**

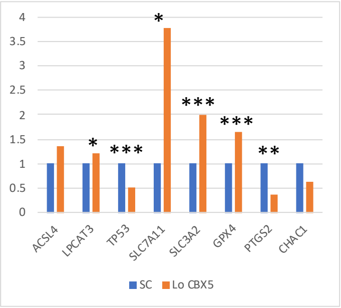


**C** Genes **down** in Low CBX5 compared to **SC**: 1350 genes

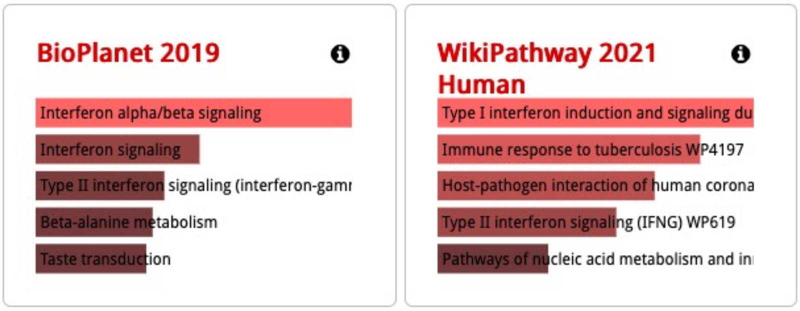


**Sup. Figure 1**

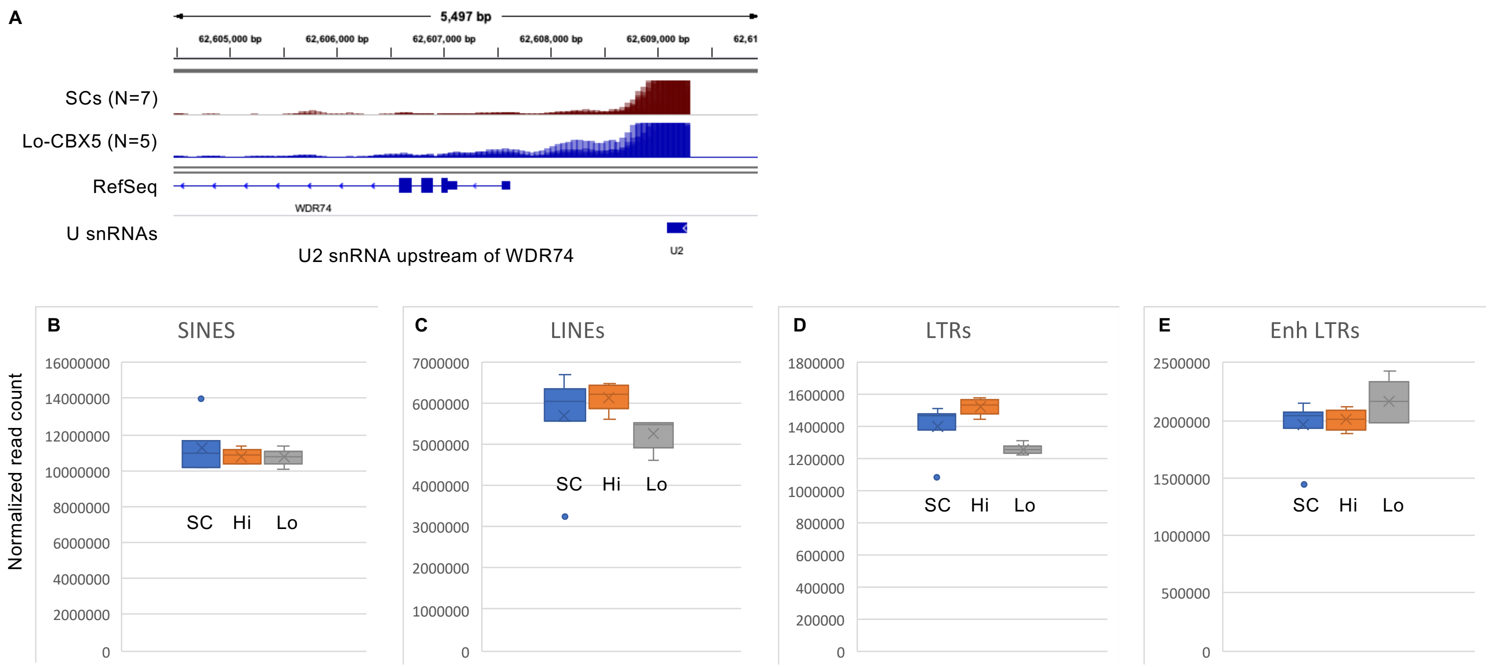
A
5,497 bp
62,605,000 bp
62,606,000 bp
62,607,000 bp
62,608,000 bp
62,609,000 bp
SCs (N=7)
Lo-CBX5 (N=5)
RefSeq
WDR74
U snRNAs
U2
U2 snRNA upstream of WDR74
B
SINES
Normalized read count
16000000
14000000
12000000
10000000
8000000
6000000
4000000
2000000
0
SC Hi Lo
C
LINEs
7000000
6000000
5000000
4000000
3000000
2000000
1000000
0
SC Hi Lo
D
LTRs
1800000
1600000
1400000
1200000
1000000
800000
600000
400000
200000
0
SC Hi Lo
E
Enh LTRs
2500000
2000000
1500000
1000000
500000
0
SC Hi Lo

**Sup. Figure 2**

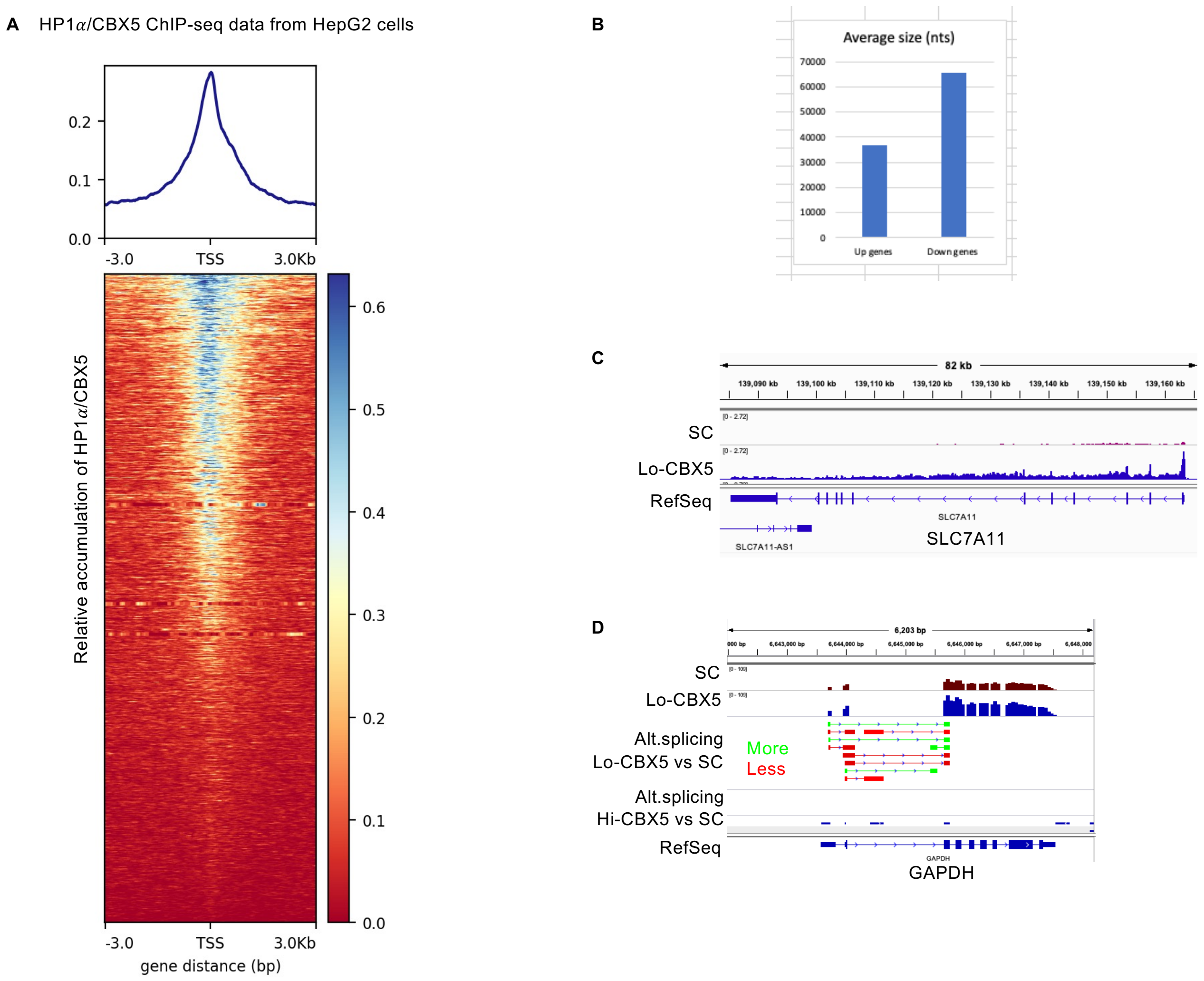
A
HP1α/CBX5 ChIP-seq data from HepG2 cells
0.2
0.1
0.0
-3.0
TSS
3.0Kb
Relative accumulation of HP1α/CBX5
0.6
0.5
0.4
0.3
0.2
0.1
0.0
gene distance (bp)
B
Average size (nts)
70000
60000
50000
40000
30000
20000
10000
0
Up genes
Down genes
C
82 kb
SC
Lo-CBX5
RefSeq
SLC7A11
SLC7A11-AS1
D
6,203 bp
SC
Lo-CBX5
Alt.splicing
Lo-CBX5 vs SC
More
Less
Alt.splicing
Hi-CBX5 vs SC
RefSeq
GAPDH

**Sup. Figure 3**

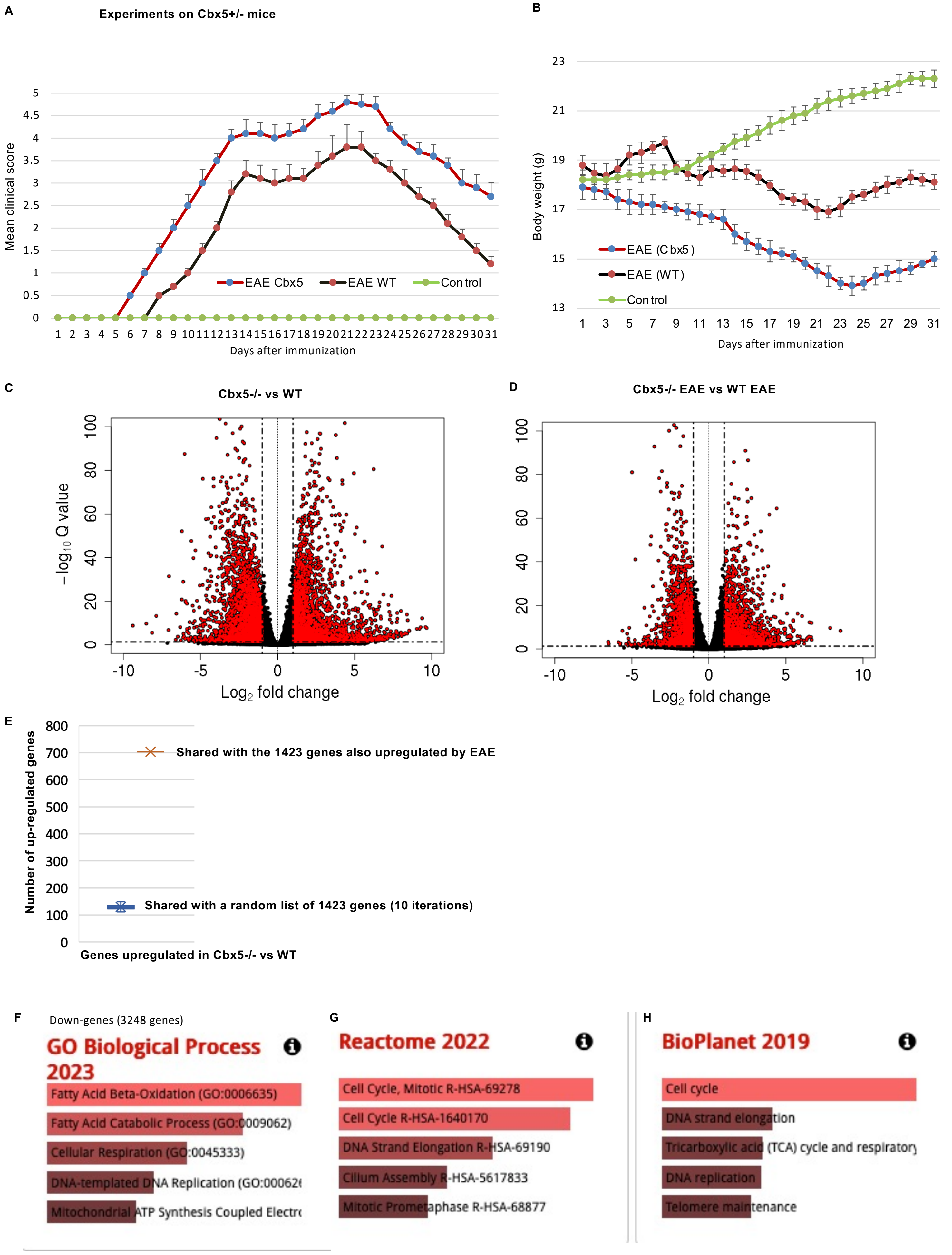
A
Experiments on Cbx5+/- mice
Mean clinical score
EAE Cbx5
EAE WT
Control
Days after immunization
B
Body weight (g)
EAE (Cbx5)
EAE (WT)
Control
Days after immunization
C
Cbx5-/- vs WT
– log10 Q value
Log2 fold change
D
Cbx5-/- EAE vs WT EAE
Log2 fold change
E
Number of up-regulated genes
Shared with the 1423 genes also upregulated by EAE
Shared with a random list of 1423 genes (10 iterations)
Genes upregulated in Cbx5-/- vs WT
F
Down-genes (3248 genes)
GO Biological Process 2023
Fatty Acid Beta-Oxidation (GO:0006635)
Fatty Acid Catabolic Process (GO:0009062)
Cellular Respiration (GO:0045333)
DNA-templated DNA Replication (GO:00626
Mitochondrial ATP Synthesis Coupled Electro
G
Reactome 2022
Cell Cycle, Mitotic R-HSA-69278
Cell Cycle R-HSA-1640170
DNA Strand Elongation R-HSA-69190
Cilium Assembly R-HSA-5617833
Mitotic Prometaphase R-HSA-68877
H
BioPlanet 2019
Cell cycle
DNA strand elongation
Tricarboxylic acid (TCA) cycle and respiratory
DNA replication
Telomere maintenance

Sup. Figure 4

A

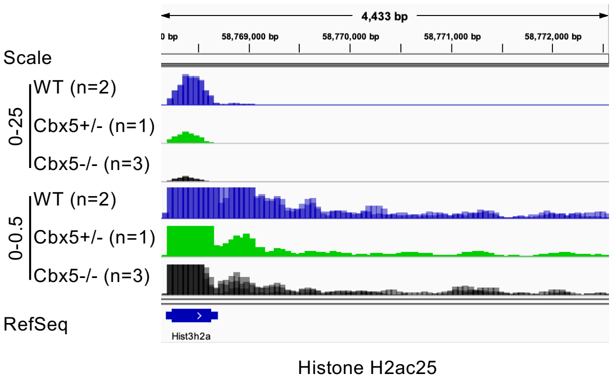


B

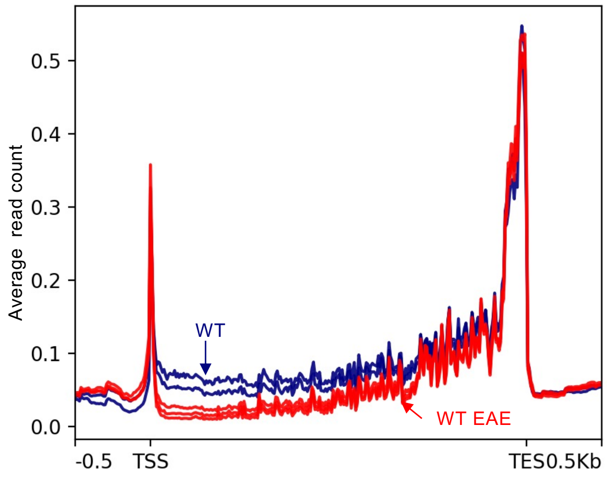


C

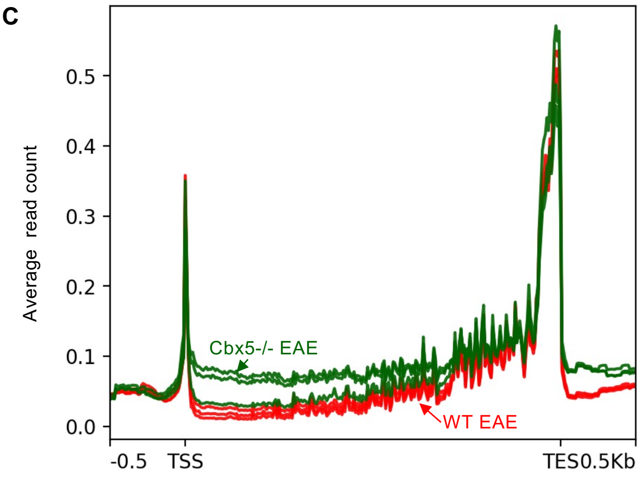


**Sup. Figure 5**
